# SNP panel for non-invasive genotyping of leopard (*Panthera pardus*)

**DOI:** 10.1101/2024.11.01.621452

**Authors:** Faruk Mamugy, Laura D. Bertola, Amber Mertens De Vry, Nicolas Dussex, Bastian Shiffthaler, Johanna L. A. Paijmans, Michael Hofreiter, Ryan Forbes, Graham I.H. Kerley, Kris Everatt, Matthew Becker, Scott Creel, Stéphanie Bourgeois, Clive Chifunte, Marine Drouilly, Göran Spong

## Abstract

Genetic resources for species monitoring should ideally be relevant for the species’ full distribution range, feasible economically and logistically, and validated for the range of sample types collected from the field. This is particularly important for large carnivores that are elusive and wide-ranging, where individual and population processes often traverse administrative borders, and where obtaining high-quality samples can be challenging. Here we present a small species-specific SNP panel for leopards. We used whole genome data from across the global range and RAD sequence data from Zambian leopards to select markers for assay development. These were ascertained for 590 individual leopards from eight African countries and final selection was based on marker variation and performance on non-invasive samples. The final 96 marker panel holds 5 mitochondrial markers for species recognition, 3 Y-markers for determination of individual sex, 3 X-markers and 85 somatic markers, with an associated genetic baseline holding nearly 900 individuals. The selected autosomal markers hold variation across the global range with high power to identify individuals (PID=2,45×10^−35^) and in most cases their provenance with high assignment probability (>95%). Markers were also selected based on their performance on samples with low target DNA content, with distinct genotype separation in the output marker plots. The genotypes from this panel are thus straightforward to analyze and do not require computationally challenging bioinformatic resources, making this a low cost and accessible resource for leopard monitoring and research.

## Introduction

Genomes hold vast amounts of information. By extracting parts of this information, individuals, their kin, reproductive patterns, behaviors, population features and history, evolutionary patterns and more can be revealed. Since their inception, genetic applications have increasingly been deployed to study wild populations and address questions difficult to assess via alternative means (REFSXX). As methods improve and costs decrease, the allure of genetic data persists. However, establishing protocols for extracting genetic data from wild populations may require a substantial development cost. Once established, the protocol must negotiate cost, logistics, and data quality and quantity to accommodate sample influx that may be constrained by time or unpredictable in size and quality.

Minute quantities of DNA, sometimes collected non-invasively from the environment, permit the extraction of complete genotypes from individuals without direct observation (Thomsen and Willerslev 2015). This possibility can be particularly useful when studying elusive species living at low population densities or to minimize disturbance to the subjects. Since such collection can be done with simple means, it greatly facilitates large-scale collection efforts by management officials or the public e.g. (Giangregorio et al. 2019; Spitzer et al. 2016; Norman and Spong 2015). While relatively easy to collect, such as in the form of fecal samples, non-invasive sample sets typically include samples at the detection threshold for molecular analysis. Such samples have low amounts of intact DNA, and this may lead to difficulties in extracting data of sufficient quality. As for all data, genetic data also hold errors that may affect downstream inferences if not appropriately controlled (Creel et al. 2003). By running samples multiple times, the prevalence and manifestation of errors can be mapped. In some cases, building consensus genotypes may be a way to generate genotypes of sufficient quality and resolution (Solberg et al. 2006; Waits, Luikart, and Taberlet 2001), but this method requires many replicates to make it statistically sound, making it demanding in resources (note that sequencing applications such as GT-sequencing in essence build a consensus genotype based on fragment prevalence). An alternative method is to use amplification success as a proxy for quality and simply discard samples below an empirically determined threshold, but for small sample sets this may not be an option. Whichever method is chosen to build genotypes, statistical validation of the final genotypes and their error rates should be performed. If genotyping error is underestimated, estimates of standing genetic variation or population size may be inflated (Creel et al. 2003). It might also cause loss of power to detect population structure and affect the precision of provenance assignments.

Any genetic marker panel should also be established for the population of interest, ideally by including samples from across the range from the very beginning of the development process. However, due to financial, logistical, and administrative constraints, such broadscale and transboundary sampling efforts remain rare, especially for wide-spread species (Kopatz et al. 2021). Yet, if the development of genetic markers only holds for a subset of the population, other subsets later included may suffer from incorrect inferences due to ascertainment bias. At its most extreme, lack of variation may prohibit identification of individuals. For species with a large, sometimes fragmented, population range, a lack of common genotype variation between subpopulations may thus limit the utility of a resource originally developed for a part of the species range (e.g. Giangregorio et al. 2019). But even when variation is sufficient for individual identification, many population genetic methods use the distribution of genetic variation to infer a range of processes such as dispersal and population size. Since such methods are generally based on the fundamental principles of population genetics (Wright 1931), false differences caused by marker ascertainment may lead to erroneous inferences, directly affecting estimates of connectivity, population size and history (Palsboll et al. 2013).

Leopards (*Panthera pardus*) have the most extensive geographic distribution of all large cats, with various subspecies ranging across a variety of habitats from the African continent and the Arabian Peninsula, to the Southeast Asian islands and as far north as Eastern Russia. Eight subspecies are recognized (Kitchener 2017, Stein et al. 2024) with all African populations considered to belong to the nominate form *P. pardus pardus*. The status of individual subspecies and populations vary dramatically: seven out of eight subspecies are threatened, with three being critically endangered, two endangered, and two vulnerable (Stein et al. 2024). Like most carnivore species, ranges have shrunk and become increasingly fragmented due to habitat loss, persecution, and illegal trade (Ripple et al 2014). Leopard populations thus currently occupy only a fraction of their historic range. Its secretive nature and adaptability has permitted the leopard to eke out a living also in areas heavily dominated by humans (Pečnerová et al. 2021), IUCN still classifies its status as ‘Vulnerable’ due to apparently declining population trends. But actual population numbers, and hence rates or extents of decline, are virtually unknown. There is therefore an urgent need for better monitoring tools across the current range to support conservation, management and law enforcement activities (Ripple et al. 2014).

Here we present a 96 marker SNP-panel optimized for non-invasive monitoring of leopards across their global range. The panel holds mt-, sex-, and nuclear markers where the vast majority show within population variation across the global range. Only markers that performed well on non-invasive material are included and the current selection provides high power to identify individuals, their sex, their kin, and their provenance.

## Materials & Methods

Our SNP-panel development leverages an existing dataset and integrates new data to improve geographic representation, optimize the statistical power, and forms the start of a baseline for analyses of provenance. First, we used the SNP variant information from previously published whole genome data of 26 individuals (including historic samples) from across the global range of leopards (Paijmans et al. 2021). Second, we identified SNPs by RAD sequencing of 152 tissue samples collected by the Zambian Carnivore Programme. These two data sets were then intersected to identify markers with variation across the global range. Third, a final marker selection step and assessment of marker performance was done on a fluid-based SNP-chip genotyping platform (Fluidigm Biomark or BioTools Juno X) using tissue and fecal extractions. Over 880 individual leopards from six African countries have now been genotyped with this panel.

### Sample collection

The data and samples included in this study come from a range of projects and initiatives. Research tissue samples were collected during ecological field studies by biopsy darting or during handling (e.g. to fit radio collars or remove wire snares). All animal handling, sample collection, and shipping were done under relevant permits renewed yearly or issued prior to sample exports. We have also included samples collected by law enforcement and organizations fighting wildlife crime. These were taken from specimens of skin, bones, and teeth known or suspected to be leopard. Scat samples were collected as per Forbes *et al*. (2024).

### Whole genome sequencing

For full details on the whole genome sequencing see (Paijmans et al. 2021). From these samples, a total of just under 1.4M SNPs were identified. In short, we used the vcf output from the sequencing to identify SNPs with global variation that overlapped with the SNPs found by the RAD sequencing below.

### Rad sequencing

Samples used for RAD sequencing (n=152) were extracted manually at the Luangwa field site of the Zambian Carnivore Programme, using the DNeasy Blood & Tissue Kit (Qiagen) or in our laboratory at SLU, Sweden by a Qiagen Symphony extraction robot using the Qiagen Tissue Extraction kit according to the manufacturer’s instructions. DNA quantity and purity were assessed by agarose gel and NanoDrop (Thermo Fisher Scientific).

We digested 0.5μg DNA with the restriction enzyme EcoRI (Thermo Fisher Scientific). Digestion quality was visualized by gel electrophoresis. We submitted samples to the National Genomics Infrastructure (NGI, SciLifeLab, Stockholm), where fragments of 400-800bp were excised for paired-end library construction (2×150bp) and RAD-sequencing on an Illumina NovaSeq 6000.

### SNP detection and filtering

Sequence quality of the demultiplexed RADseq Illumina reads was assessed using FastQC (version, ref.). Raw reads were trimmed and adapters removed with Trimmomatic (0.39, ref.) in paired-end mode. Only reads with length >100 bp after trimming were retained for further processing. These remaining sequences were used as input for SNP detection using Stacks (2.53, ref.), running the individual components of the pipeline manually. A total of 6,931 SNPs were identified using the Stacks software. We filtered SNPs using customized R scripts to remove those of low quality. First, we selected only bi-allelic SNPs that were represented in at least 75% of the samples. Only stacks with one SNP with a minimum of 30bp up- and downstream (for assay development) were retained. with a minor allele frequency (MAF) > 0.2 and X-chromosome SNPs with MAF > 0.15. Finally, we selected for SNPs in which at least 5 samples were attributed to each genotype (i.e. aa, bb and ab) and that showed variation across the global range. After applying these filtering criteria, 148 SNPs were selected for assay design. A total of 133 SNPs passed the *in silico* assay design, and were brought forward to *in vitro* assay development on a Fluidigm Biomark or Juno X.

Published Genbank sequences were used to identify Y-chromosome SNP markers that are informative of parental lineage (Table 1) and sex (Table 2). The selected sequences were aligned using BioEdit (v. 7.2, Tom Hall, Ibis Biosciences, Carlsbad, USA) and manually scanned to identify SNPs. The three sex-specific Y-chromosome SNPs simply detect the presence of the Y-chromosome (holding no variation). When these markers fail to amplify, samples are called female, and vice versa. Additionally, heterozygosity in the X-chromosome markers, only possible in females, is also used to verify sex assignments when possible. Mitochondrial SNPs were included to study matrilines and to distinguish from other sympatric carnivore species, such as lion (*Panthera leo*) and cheetah (*Acinonyx jubatus*), particularly relevant for fecal species identification. Regions with known numts were excluded.

**Table 1.** Summary marker statistics. H_o_-Observed heterozygosity, H_T_-Expected heterozygosity, F_ST_-Standardized fixation index, MAF-Minor Allele frequency.

| Marker type (N) | $H_O$ | $H_E$ | $F_{STP}$ | MAF |
| --- | --- | --- | --- | --- |
| mitochondrial | N/A | N/A | N/A | N/A |
| Y | N/A | N/A | N/A | 0.25 |
| X | 0.22 | 0.16 | - | 0.09 |
| Autosomal | 0.58 | 0.56 | 0.02 | 0.17 |

### SNP validation

The autosomal and X-chromosome SNP markers were initially validated through genotyping a total of 119 leopard samples (2 lions and 1 cheetah) using the Fluidigm Biomark platform. The mitochondrial and Y-chromosome markers were validated by genotyping 31 leopards, (59 lions and 1 cheetah,) including 9 leopards of known sex (female n = 5, male n = 4). Both positive (sample replicates) and negative no template controls were included. We evaluated the call rate of the DNA samples and the performance of the assayed markers at successfully assigning a genotype to the samples. Every marker was visually assessed on the allele clustering of the DNA samples in the Biomark scatterplot. SNPs were invalidated if the controls did not work as expected, if the different genotypes clustered too near each other or if not all genotypes were present (i.e., XX, XY, YY). Only SNPs that passed these criteria were included on the final SNP-chip (with the exception of mitochondrial and Y-markers).

The selected final SNPs (N=96) were used for a final genotyping run of sequenced individuals and additional leopard samples across Africa. Samples were collected from different countries, covering broader variation in the African leopard (Pečnerová et al. 2021), as shown in Figure 1, and were extracted from a range of source material (blood, feces, hair, and tissue). Sample replicates and descriptive statistics, such as heterozygosity and amplification success, were used to quality control data. Prior to assignment analyses, we removed replicates of individuals (individuals sampled more than once) by running the package Allelematch. We also removed samples with low amplification rate (more than 5% missing data), resulting in a final set of 875 individuals prior to the assignment analysis. To test the SNPs performance as markers for investigations of population structure, we conducted a Discriminant Analysis of Principal Components (DAPC) analysis using the Adegenet package in R 4.3.2. We analyzed structure without *a priori* spatial information to explore the number of genetic groups in the data. Individuals were then assigned to the identified genetic groups.

**Figure 1.**
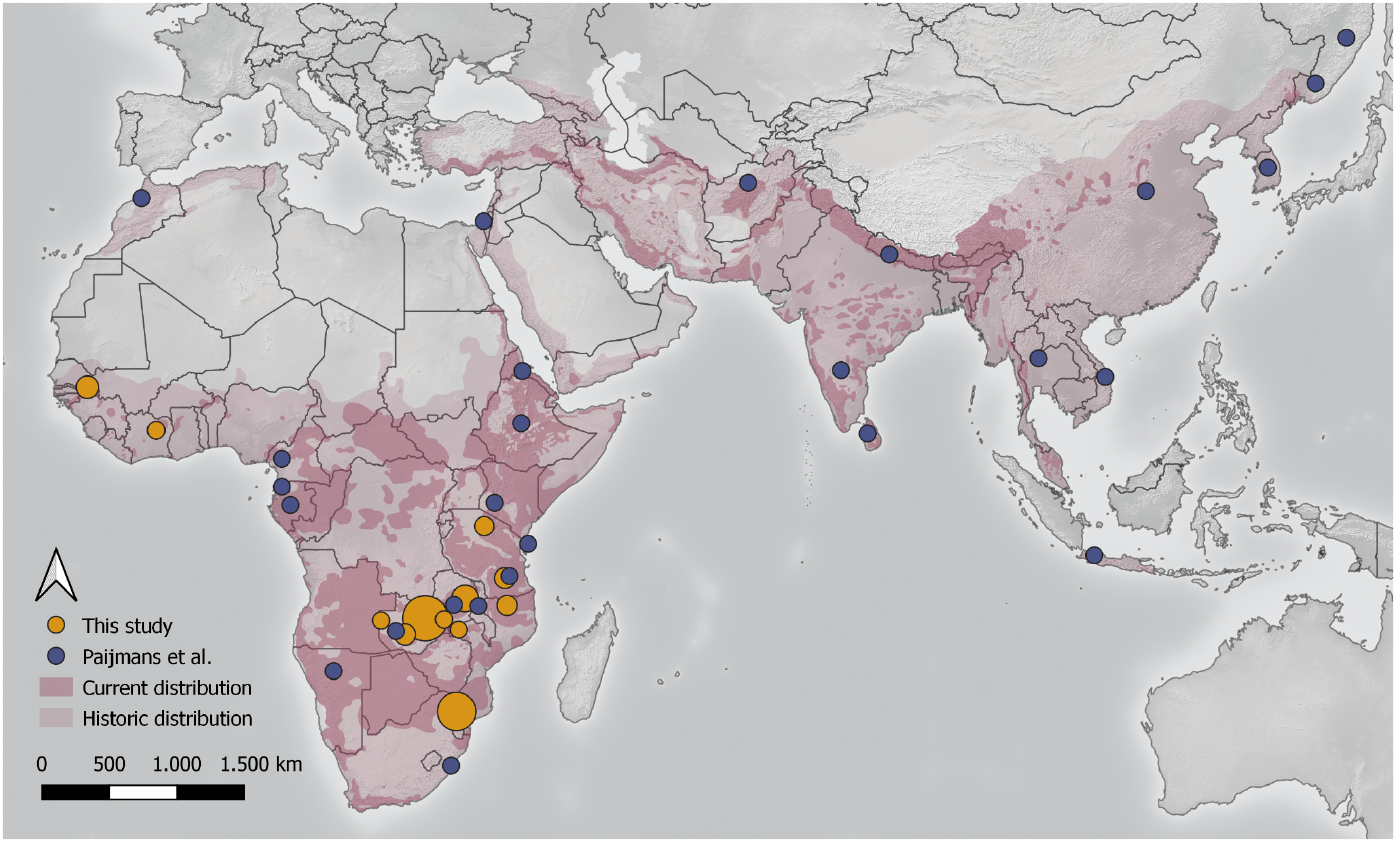
Sample localities for leopard samples used in the design and testing of the SNP panel. Blue sample locations indicate whole-genome sequencing data (1 sample per locality), orange sample localities indicate newly integrated data (larger circles represent a higher number of samples - see Suppl. Table X). Leopard distribution derived from the IUCN Red List (Stein et al. 2024), with current distribution using both ‘resident’ categories (‘extant’ and ‘possibly extant’).

### SNP performance

We assessed the reliability of genotype calls by running samples in replicates (N=39). We replicated both tissue and fecal extracts. Unfortunately we did not have access to both tissue and fecal samples from known individuals, but instead compared the distribution of individual allelic differences to the distribution of errors (Figure 2). This allowed us to empirically determine an amplification threshold (>98%) that suppresses the error rate to a point where the appearance of ghost or shadow genotypes is extremely unlikely (p<0.001).

**Figure 2.**
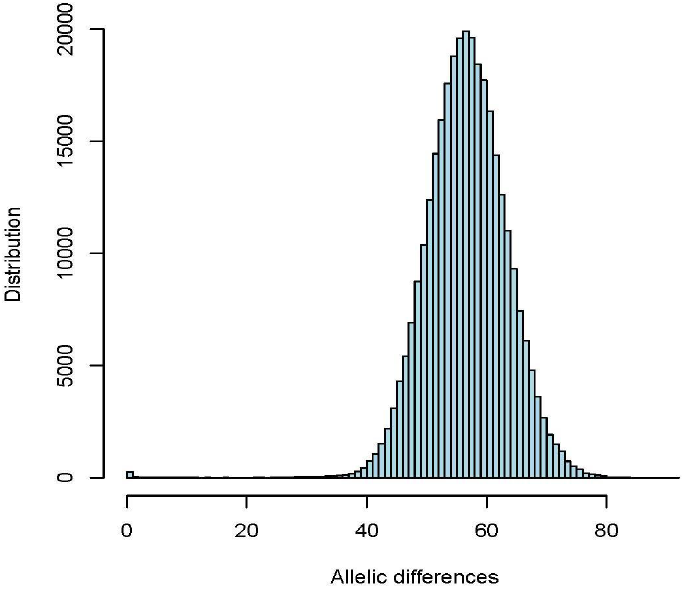
Pairwise allelic differences between all sample pairs. The large peak on the right, shows the distribution of genotype differences between unique individuals. The small peak on the left, shows discrepancies between sample pairs originating from the same individual, where the manifested genotyping error rate is 3% when excluding samples with an amplification rate <98% (for samples with an amplification rate as low as 90% the error rate is <0.06%). With lower amplification rates, sample quality may become prohibitive for reliable genotyping risking ghost and/or shadow effects in the binning of genotypes.

## Results

When samples hold enough DNA to result in high amplification rates, the individual genotypes provide reliable individual identification. For genotypes that passed the quality control filter, the mean missing data were <0.015.

The sex-determination markers correctly assigned all samples of known sex, resulting in an error rate of zero. Our Y markers hold no variation, but are included with the sole purpose of identifying sex. In contrast the X markers do hold variation, so in addition to providing additional individual and population resolution, the X markers can be used to further indicate sex-Males should appear homozygote for both Y and X, whereas females may be heterozygote for X (and lack any amplification for Y). For summary marker statistics, see Table 1.

Our quality control protocols provided empirical evidence that genotyping error can be controlled for successful individual identification (Figure 2), also from non-invasively collected samples. Even samples stored under suboptimal conditions were used to extract genotypes of sufficient quality. This is a critical requirement for most applications, but in particular when individual identity is of paramount importance (e.g. Capture-Mark-Recapture or forensics).

Based on the SNP genotyped samples, spanning a significant part of the African leopard distribution, three groups were identified, one holding west African samples, one holding mostly samples from South Africa and Mozambique (Greater Limpopo Area), and one large group holding samples from Tanzania, Zambia and northern Mozambique. When re-analysing the two latter groups (i.e., excluding west African samples), this group split into three genetic groups (Figure 3b and Table S2).

**Figure 3.**
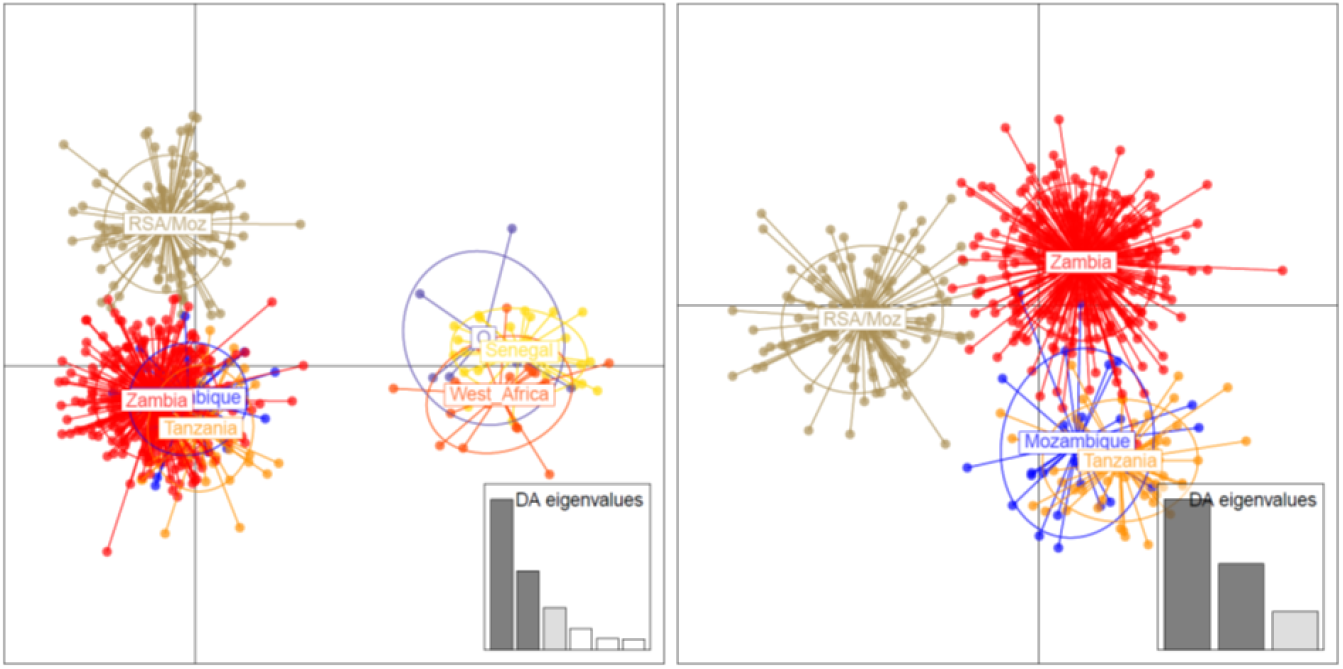
Discriminatory Analysis of Principal Components (DAPC). a) DAPC of all samples showing clear distinction between samples from west Africa. b) Subset of samples to only include East and southern Africa. Samples separate into groups with high alignment to geographical origin of samples (Table S2).

## Discussion

Methods to extract genetic data from increasingly degraded sources are rapidly evolving and costs are declining. For applications across research, management and law enforcement these advances hold much promise. But moving from possibilities to practicalities when acquiring genetic data will invariably be a trade-off between **cost, genotype quality** and **logistics**. The optimal solution is also likely to change over time as methods improve and projects evolve. In this study we describe how we have resolved this trade-off to build a resource for research and conservation of leopards, applicable throughout the global range and optimized for low quality samples.

To save on **costs**, existing genomic resources, e.g. full genomes derived from previously published studies, can be repurposed to design cost-effective tools, which can meet the demand for cheap, accessible and scalable tools. Here we have presented a genotyping panel for leopards optimized for samples of low quality, partially dependent on previously published whole genome data and new RAD sequencing data. For most queries of management and law enforcement, including many questions in behavioral ecology, the ability to identify individuals, their sex, their kin, population structure and provenance reliably is what is needed. This allows for robust population estimates through capture-mark-recapture, estimates of connectivity, parentage analyses, assessments of within-population diversity, and assignment of unknown samples to a region of origin. While additional markers would likely further increase the resolution for more demanding analyses such as pedigree reconstruction and provenance, it would also raise costs of development and downstream genotyping. Once a set of markers has been developed, the cost of genotyping is to a large extent dependent on scale. If samples cannot be run in suitable batch sizes, the cost may quickly become prohibitive. Alternatively, delaying analyses until a sufficient sample number has been accrued may delay results that may urgently be needed. Here, the ability to run a smaller set of samples is useful, for example by using single tubes in a qPCR or systems that allow flexible plate configurations (e.g. Biosystems Juno).

Our **quality control** results align well with our previous work on other model systems (Creel et al. 2019; Kleven et al. 2019; Norman and Spong 2015; Walton et al. 2021) and other studies showing that SNPs are better suited for working with degraded DNA than microsatellites (Morin et al. 2004). Additionally, liquid-based SNP arrays have shown to be better suited for low DNA content samples, than the more commonly used prespotted arrays (REFXX). During the optimization of this panel, we included scats collected along the South Africa/Mozambique border > 8 years prior to DNA extraction and genotyping, with the dried scats stored in doubled air-tight plastic bags with silica gel in the inner bag. Additionally, temperatures in the storeroom in which the scats were stored regularly exceeded 40 °C during summer. Despite the suboptimal conditions for DNA preservation, high-quality genotypes were successfully acquired from 52% of the scats identified as leopards in the field. The high performance of our SNP panel with degraded DNA may enable researchers to proactively and opportunistically collect samples and store them, while for example, funding is acquired, and still obtain high genotyping success rates.

The **logistical** challenges of working with wild populations, often of conservation concern, can be ameliorated by the protocols chosen. *First*, the inherent qualitative nature of SNPs allows for reliable and direct replication across laboratories and platforms. As such, and although whole-genome sequencing (WGS) is dropping in price rapidly, genotyping a preselected SNP panel is still a cost-effective and pragmatic tool for monitoring wildlife for the foreseeable future. Many questions of interest to wildlife managers do not require full genomes or a high number of SNP markers, but can be appropriately addressed by a panel of a few dozens of SNPs. *Second*, liquid based SNP platforms are especially suitable for working with low quality samples (Carroll et al., 2018; A. von Thaden et al., 2017), such as non-invasively collected fecal samples, forensic traces, and historic samples. In research and management of wide ranging and elusive species, non-invasive samples are often the preferred or only available DNA source. Non-invasive sampling may also allow, in addition to minimizing disturbance to the animals, estimates of population size in a Capture-Mark-Recapture framework (Spitzer et al. 2016). *Third*, the ability to reliably genotype individuals from non-invasively collected material also greatly facilitate transport logistics since these samples or DNA extracts are [exempt from CITES](https://cites.org/sites/default/files/eng/cop/16/doc/E-CoP16-25-A3.pdf). This significantly reduces the time from collection effort to result. In contrast, moving samples that fall under CITES are contingent on clearance procedures that sometimes can be slow or held up by administrative differences in how these co-dependent import- and export-permits are issued. Moreover, CITES permits are only valid for a specified number of samples, making it virtually impossible to apply for a permit before the collection effort has been completed. *Finally*, an open source SNP panel for the full distribution of the species, may ameliorate the imbalance in international collaborations by reducing the dependence of colleagues in range states on sequencing and computational facilities in the global north. Due to their modular character, SNP panels can be set up to address different relevant population genetics questions (Alina von Thaden et al., 2020), in manners compatible with locally available equipment and genotyping platforms (e.g. desktop sequencing or qPCR). Data storage and analysis do not require advanced computer clusters, and are more accessible for those without advanced bioinformatics training.

## Acknowledgements

This work has received financial support from NSF, Vr (Swedish Research Council), SIDA, USFWS, and Panthera. We also acknowledge sequencing support from the Swedish National Genomics Infrastructure (NGI) at the Science for Life Laboratory in Uppsala and Stockholm supported by the Swedish Research Council and the Knut and Alice Wallenberg Foundation.

## Supplementary material

**Table S1.**
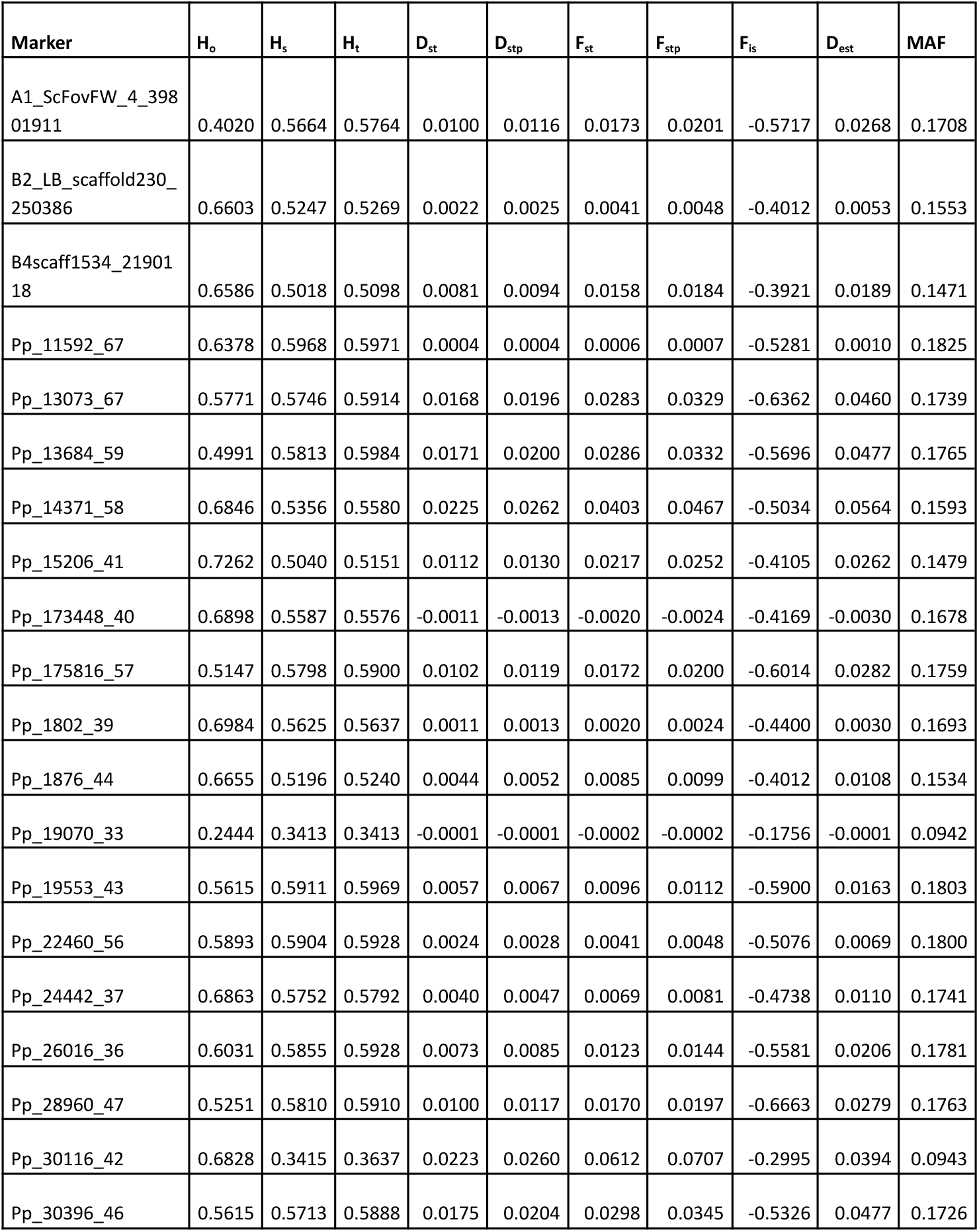
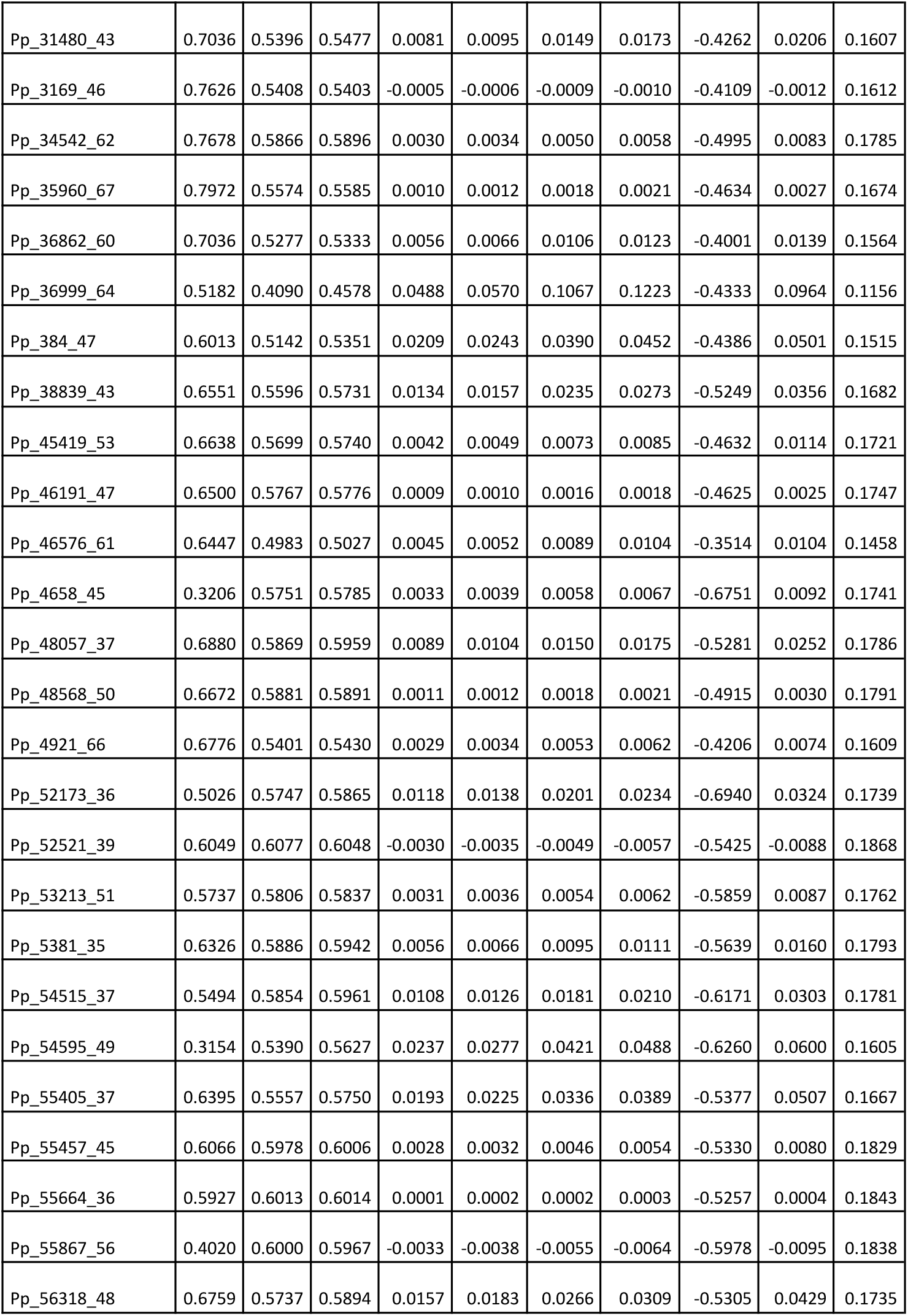
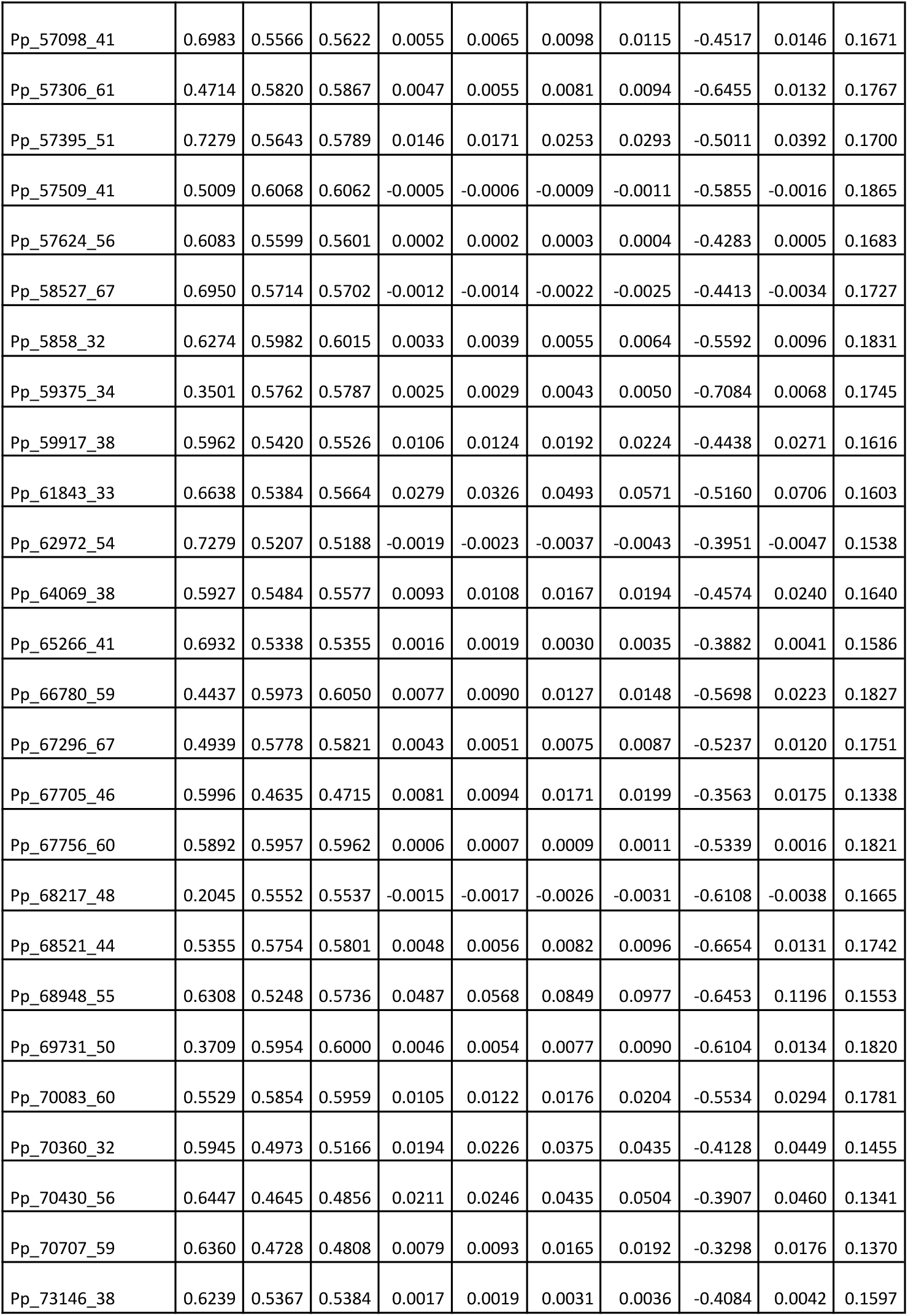
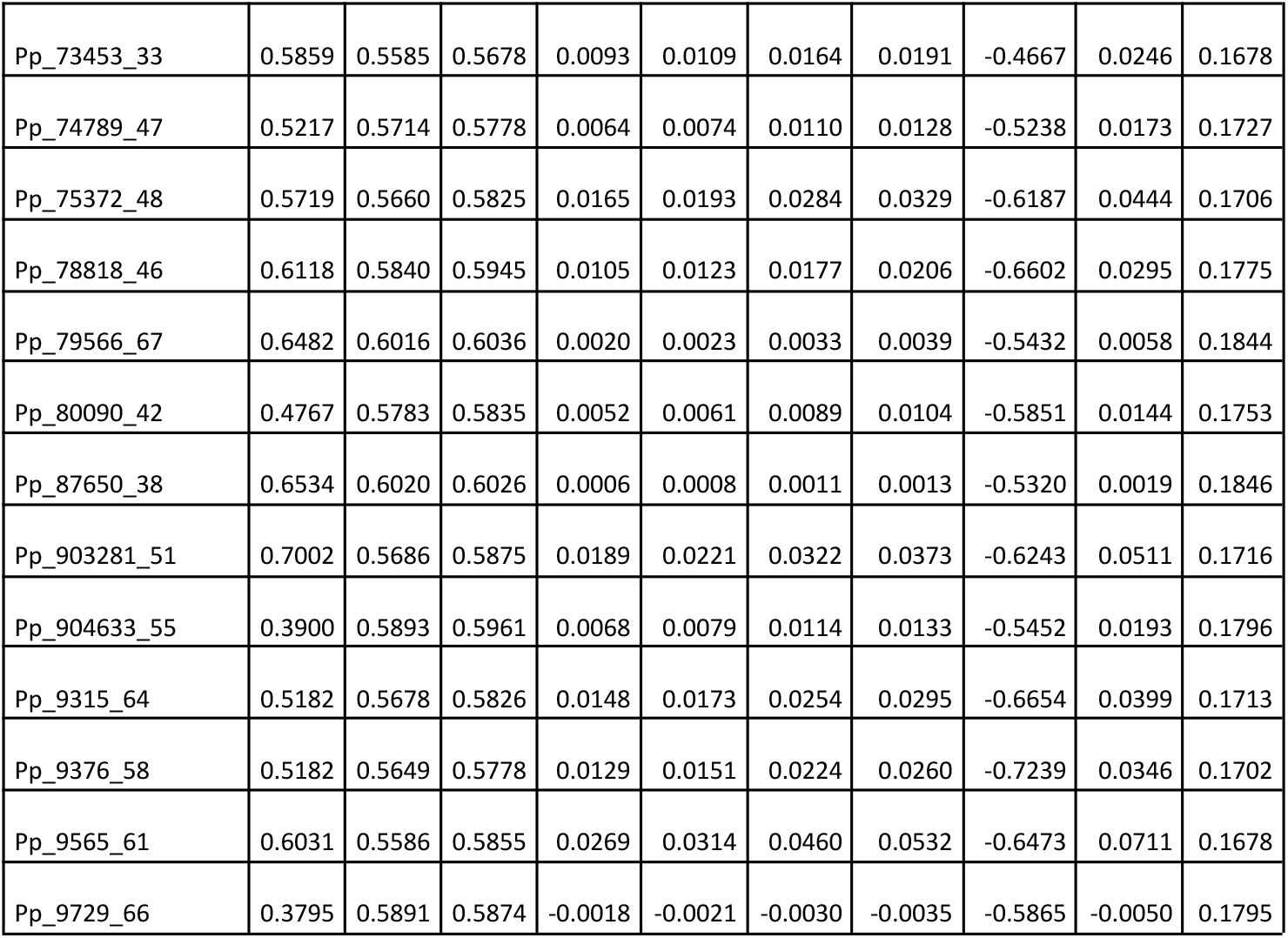
Somatic marker statistics as calculated by the R package Adegenet. Ho: Observed heterozygosity, H_s_: Expected heterozygosity within subpopulations, H_t_: Total expected heterozygosity, D_st_: Genetic diversity between subpopulations, D_stp_: Proportion of genetic diversity between subpopulations, F_stp_: Standardized fixation index, F_is_: Inbreeding coefficient within subpopulations, D_est_: Jost’s measure of differentiation, MAF: Minor allele frequency.

**Table S2.**
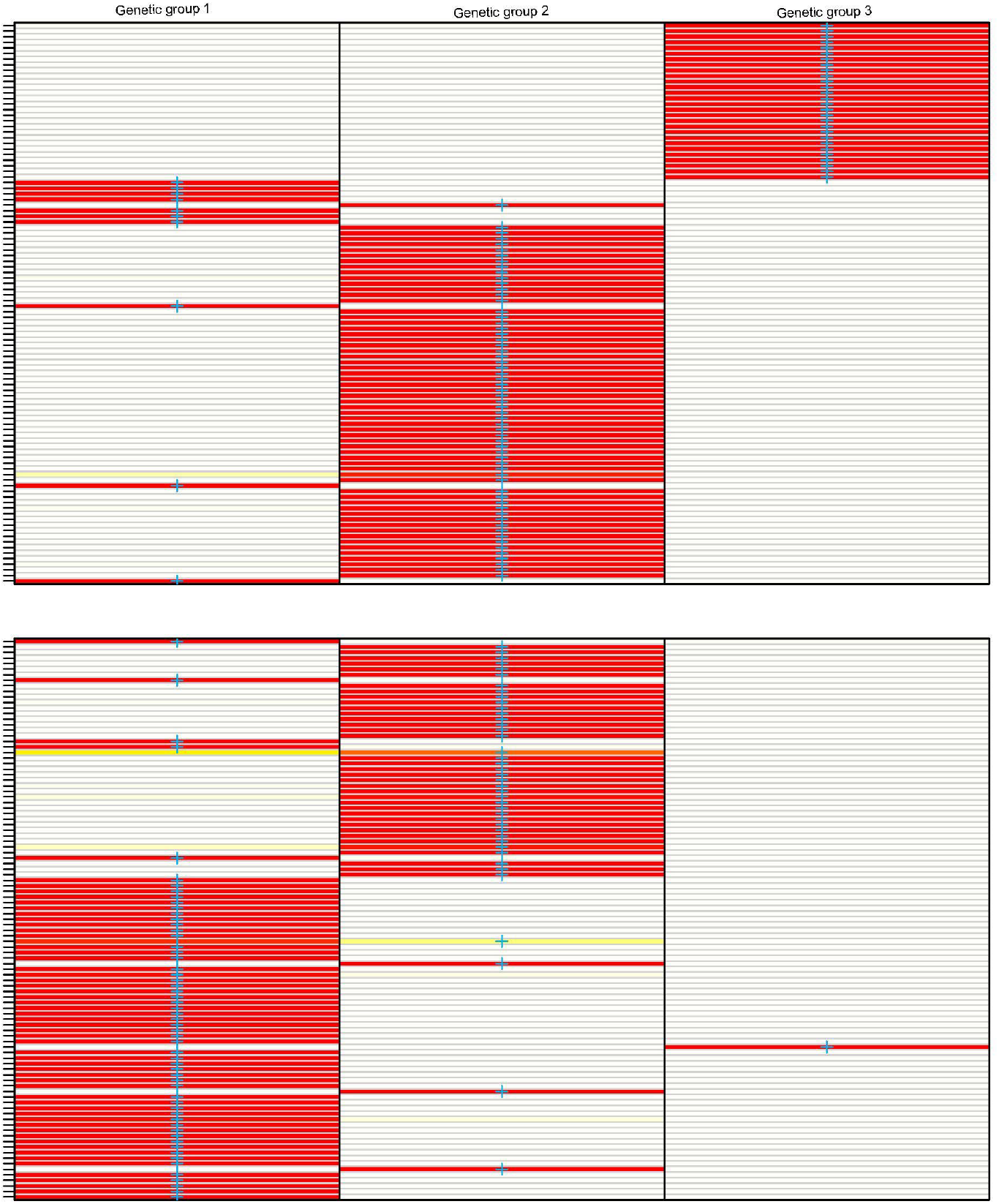
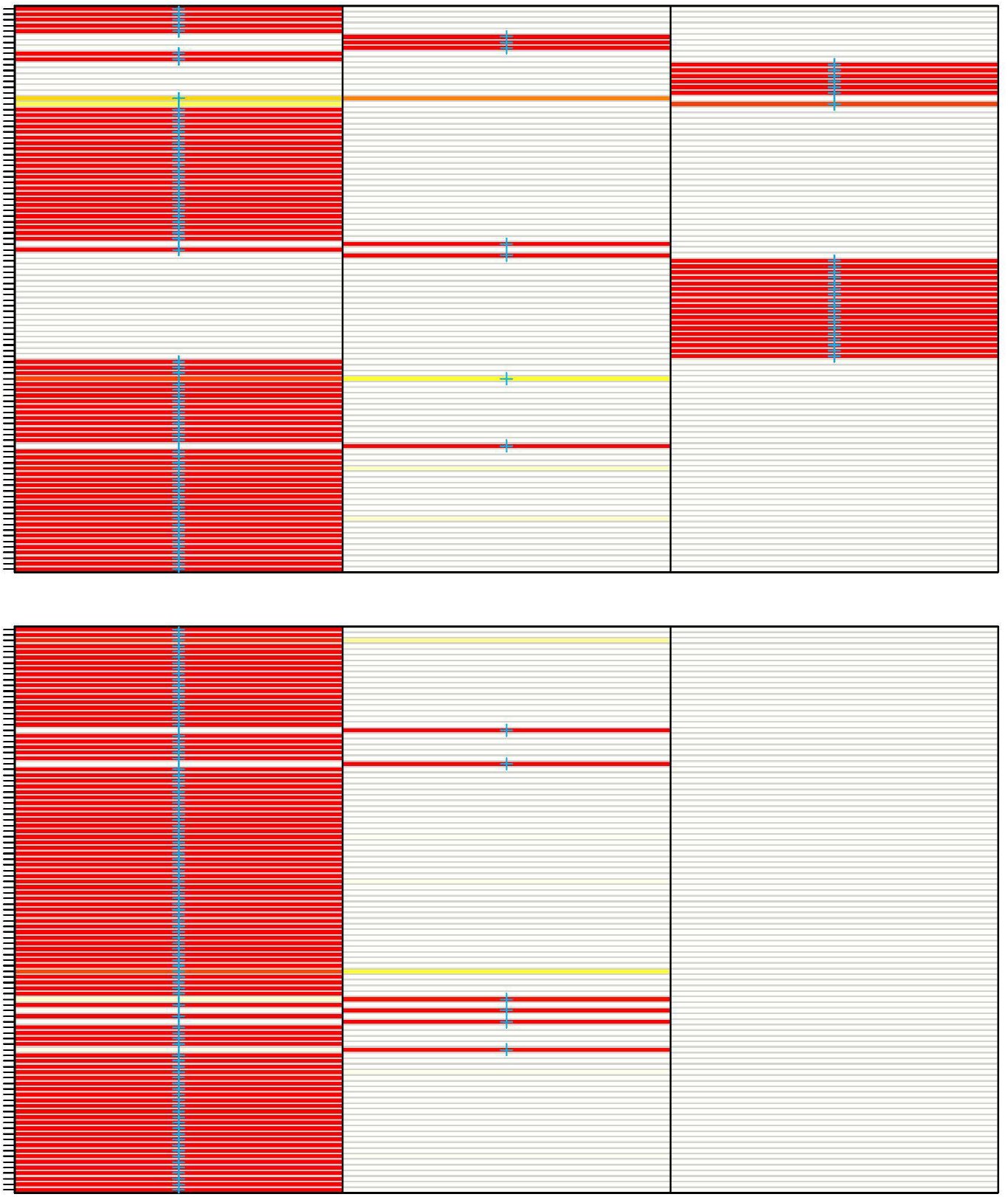
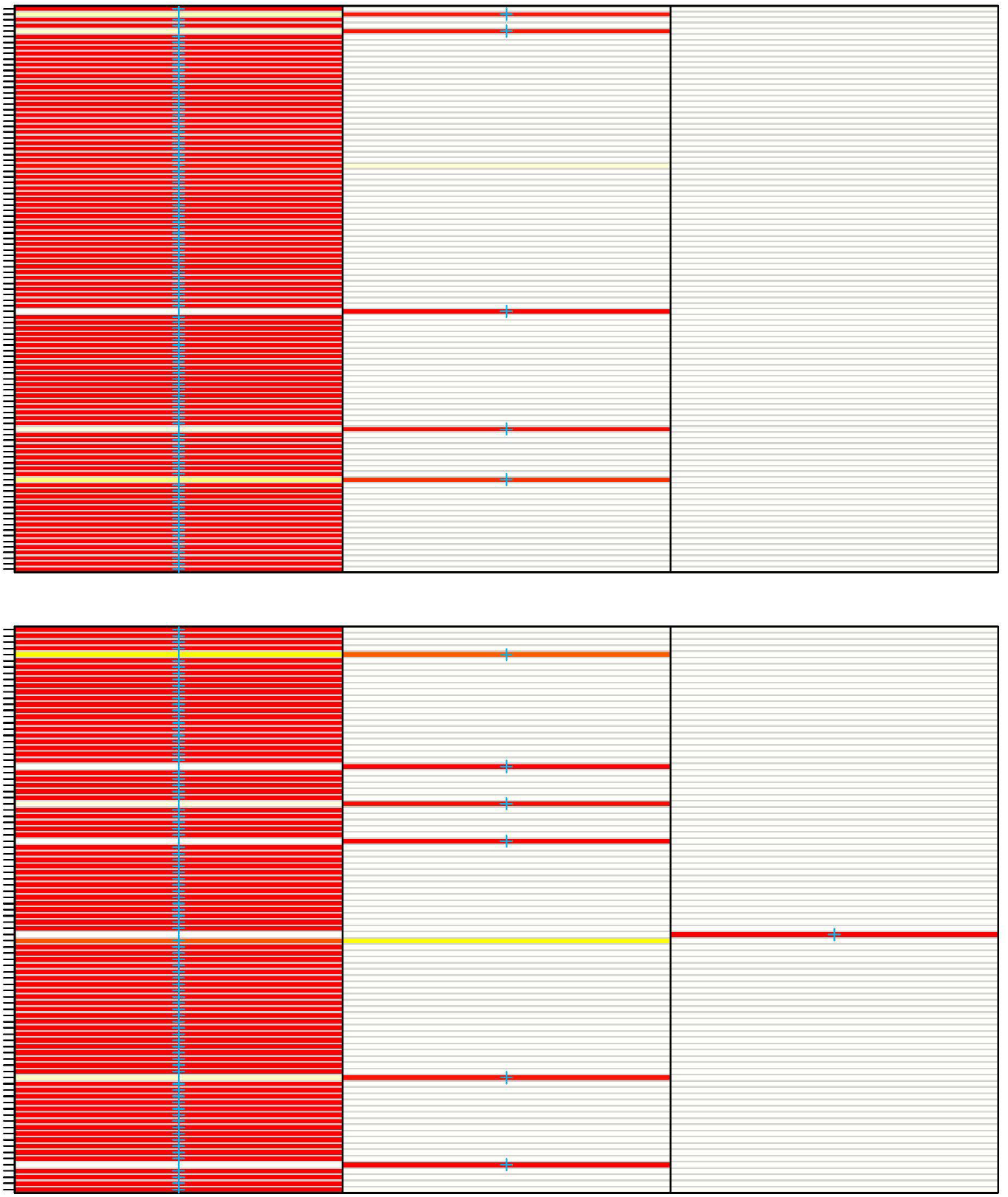
Individual assignment probabilities to the three genetic groups shown in figure 3b. Assignment probabilities >95% are colored in red.

